# Molecular dietary analyses of western capercaillies (*Tetrao urogallus*) reveal a diverse diet

**DOI:** 10.1101/2021.03.08.434346

**Authors:** Physilia Ying Shi Chua, Youri Lammers, Emmanuel Menoni, Torbjørn Ekrem, Kristine Bohmann, Sanne Boessenkool, Inger Greve Alsos

## Abstract

Conservation strategies centred around species habitat protection rely on species’ dietary information. One species at the focal point of conservation efforts is the herbivorous grouse, the western capercaillie (*Tetrao urogallus*). Traditional microhistological analysis of crop contents or faeces and/or direct observations are time-consuming and at low taxonomic resolution. Thus, limited knowledge on diet is hampering conservation efforts. Here we use non-invasive environmental DNA (eDNA) metabarcoding on DNA extracted from faeces to present the first large-scale molecular dietary analysis of capercaillies. Faecal samples were collected from seven populations located in Norway (Finnmark, Troms, Trøndelag, Innlandet) and France (Vosges, Jura, Pyrenees) (n=172). We detected 122 plant taxa belonging to 46 plant families of which 37.7% of the detected taxa could be identified at species level. The average dietary richness of each sample was 7 ± 5 SD taxa. The most frequently occurring plant groups with the highest relative read abundance (RRA) were trees and dwarf shrubs, in particular, *Pinus* and *Vaccinium myrtillus*, respectively. There was a difference in dietary composition (RRA) between samples collected from the different locations (adonis F_5,86_= 11.01, p <0.05) and seasons (adonis F_2,03_= 0.64, p <0.05). Dietary composition also differed between sexes at each location (adonis F_1,47_ = 2.77, p <0.05), although not significant for all data combined. In total, 35 taxa (36.84% of taxa recorded) were new capercaillie food items compared to existing knowledge. The non-invasive molecular dietary analysis applied in this study provides new ecological understanding of capercaillies’ diet which can have real conservation implications. The broad variety of diet items indicates that vegetation does not limit food intake. This plasticity in diet suggests that other factors including disturbed mating grounds and not diet could be the main threat to their survival.

## INTRODUCTION

Over the past few centuries, anthropogenic actions such as urbanisation, habitat fragmentation, over-harvesting and introduction of invasive species have accelerated the loss of biodiversity worldwide (Ceballos et al., 2010; Pimm, 2001). As we enter the sixth mass extinction, there is a critical need to obtain ecological data, which can be used to alleviate the decline of species through conservation strategies (Ceballos et al., 2015; Cracraft, 1995). To formulate effective conservation strategies, reconstruction of animal diet plays an important role in understanding species interactions, ecosystem functions and habitat selection (Clare, 2014; Kunz et al., 2011). As diet is intrinsically linked to habitat selection, diet information is a key element used for planning appropriate habitat protection strategies (Gonzalez et al., 2012), and for managing habitat restoration projects (Iwanowicz et al., 2016). In herbivores, diet studies have been widely applied in ecological studies including temporal variability in diet compositions (Aziz et al., 2017; Rayé et al., 2011), foraging plasticity (Kowalczyk et al., 2019; Quéméré et al., 2013), resource competition and niche partitioning (Kartzinel et al., 2015; Lopes et al., 2015; Soininen et al., 2015; ter Schure et al., 2020), impacts of herbivores on vegetation (Bilodeau et al., 2014; Hibert et al., 2011), and diet authentication of grazing livestock (Pegard et al., 2009).

The herbivorous grouse species, the western capercaillie (*Tetrao urogallus*), has been the focus of conservation efforts centred around boreal forest habitat protection and restoration (Ilse Storch, 2007). Due to its strong association with mature old-growth forests and its relatively large home range, the western capercaillie is also an indicator species for forest biodiversity conservation (Suter et al., 2002). Capercaillies are widely distributed across the boreal and montane forests in Eurasia, but these birds are particularly susceptible to anthropogenic disturbances, including land-use change, habitat destruction, and forest management practices (Klaus & Bergmann, 1994; Ilse Storch, 2007). As a result, populations in Central and Western Europe are fragmented due to habitat loss and uneven distribution patterns of conifer forests, and local extinctions have been recorded in the French Alps and Britain (Duriez et al., 2007; I. Storch, 2000). Capercaillies are now listed in Annex I of the Birds Directive in European countries, where suitable habitats must be protected to ensure their survivability. To obtain a complete understanding of the capercaillie habitat, detailed knowledge of its diet and how this varies throughout the species’ range is a crucial step for conservation management.

Capercaillies are mainly folivores, and they spend a considerable amount of time foraging due to low intake rate (Blanco-Fontao et al., 2010; Sedinger, 1997). Specialisation on food resources depends on local conditions, where only a few plants are important diet items throughout the capercaillies’ range (Sedinger, 1997). Traditionally, capercaillie diet studies were based on direct observations during foraging events (Gustafsson, 2008), but capercaillies are highly susceptible to human disturbances making direct observations challenging and detrimental to their well-being (Duriez et al., 2007; Mikoláš et al., 2015). Another method commonly used is the microhistological identification of plants through analysis of crop contents (Borchtchevski, 2009; Wegge & Kastdalen, 2008). This method is invasive, requiring capercaillies to be hunted and killed. Other non-invasive means have also been employed such as browsing signs analysis (Gustafsson, 2008) and microhistological identification of ingested plants in their faecal samples (Blanco-Fontao et al., 2010; Gonzalez et al., 2012; Odden et al., 2003; N. Picozzi et al., 1996; Nicholas Picozzi et al., 1999; Rodríguez & Obeso, 2000). However, these non-molecular methods are extremely time-consuming, requiring trained researchers to accurately identify their diet, and identifications are prone to errors or observer bias (Sheppard & Harwood, 2005). Easily digested plants often go undetected, and detected plants are often identified at a low taxonomic resolution (Shrestha & Wegge, 2006). The diet identified using such methods is therefore typically described using only common names such as ferns, grasses, heaths and mosses, without species or even genus-specific resolution (Blanco-Fontao et al., 2010; Gonzalez et al., 2012; Odden et al., 2003). With the advent of high-throughput sequencing (HTS) technologies such as DNA metabarcoding, an alternative non-invasive mean of studying their diet through faecal samples is possible (Valentini, Miquel, et al., 2009). This reduces the effect of human disturbances on the capercaillies as faecal samples can be readily collected after key foraging periods, to decrease the risk of flight or altered behaviour (Thiel et al., 2008).

DNA metabarcoding of faecal samples to characterise diet has been used in many herbivore diet studies (Erickson et al., 2017; Iwanowicz et al., 2016; Kowalczyk et al., 2019; Soininen et al., 2015; Valentini, Miquel, et al., 2009). It is a HTS technique that amplifies taxonomically informative genetic regions in complex or environmental samples such as faeces, by targeted PCR amplification using generic primers (Valentini, Miquel, et al., 2009). Metabarcoding allows for large-scale parallel sequencing of samples from many individuals (Valentini, Miquel, et al., 2009), which is cost-effective and allows for population-scale diet studies (Quéméré et al., 2013). Compared to traditional methods, metabarcoding can give better taxonomic resolution and identify a wider breadth of diet items provided that a well-populated species reference database is available (Soininen et al., 2009; Srivathsan et al., 2016; Valentini, Pompanon, et al., 2009). Metabarcoding can also be used to reconstruct diet in degraded faecal samples not collected immediately after defecation (Hawlitschek et al., 2018). Such flexibility is advantageous when studying the diet of elusive animals like the capercaillies, where care should be taken not to disturb them, particularly during foraging periods and the breeding season.

Currently, only one study has used metabarcoding to reconstruct the diet of capercaillies, and this study included less than ten individuals (Valentini, Miquel, et al., 2009). As such, the potential for large-scale metabarcoding analysis of capercaillie diet to retrieve a wider breadth of diet items has not been explored. Here, we present the first large-scale metabarcoding diet analysis of the western capercaillies and by doing so, we can obtain more detailed diet information across the different spatial and temporal scales. This information can be used to develop appropriate conservation measures to ensure the survival of the capercaillies. We aim to 1) document the dietary richness and composition of the plants consumed by capercaillies, 2) explore diet variation between capercaillie populations across Norway and France, 3) explore the diet variation between sexes and seasons, and 4) assess whether metabarcoding retrieves any new plant diet items as compared to previously recorded plant diet items obtained through traditional non-molecular methods. To achieve these aims we carry out metabarcoding of western capercaillie faecal droppings collected in spring and autumn from populations in seven different locations across Norway and France (Norway: Finnmark, Troms, Trøndelag, Innlandet. France: Vosges, Jura, Pyrenees). These populations are representative of the three major geographical regions inhabited by the capercaillies; Northern Europe (Norway), Central Europe (Vosges and Jura), and Southern Europe (Pyrenees, one of the southernmost capercaillie populations in the world). Hence, this study could reveal insights into the capercaillie’s diet across its geographical range.

## MATERIALS AND METHODS

### Study sites and faecal collection

In Norway, capercaillies are classified as Least Concerned (LC) and can be found in the boreal forests across most of the country (https://www.biodiversity.no/). The study areas are mostly dominated by pine (*Pinus sylvestris*), spruce (*Picea abies*), birch (*Betula pubescens*), aspen (*Populus tremula*), and ericaceous shrubs (Wegge & Kastdalen, 2008). The estimated population size is around 25,000 to 30,000 individuals (http://www.birdlife.org). In France, capercaillies are highly threatened and mostly restricted to the Vosges, Jura, and Pyrenees mountains, with an estimated population size of less than 50, 300, and 4000 individuals respectively (http://www.birdlife.org; Parc Naturel Régional des Ballon des Vosges, 2019; Observatoire des Galliformes de Montagne, 2020). They are classified as vulnerable (VU) in Pyrenees, and endangered (EN) in both Vosges and Jura (https://uicn.fr/listes-rouges-regionales/). The study area in Vosges and Jura are dominated by spruce (*Picea abies*), fir (*Abies alba*) and beech (*Fagus sylvatica*). For Vosges, there is a fairly good diversity of both coniferous and deciduous trees on the edges of clearings, and peat bogs can be found in the forests. Depending on the soil and forest cover, rich herbaceous vegetation or ericaceous field layer is present. For Jura, higher elevation parts are dominated by subalpine mountain pine woods (*Pinus uncinata*) with rich herbaceous vegetation on limestones. However, ericaceous plants are locally present on decarbonized humus. In the Pyrenees, the mountain forests are strongly dominated by fir (*Abies alba*), beech (*Fagus sylvatica*) or pine (*Pinus uncinata*). Near the tree line, there is a rich mixture of different species of trees such as rowan (*Sorbus aucuparia*) and birch (*Betula verrucosa*), shrublands dominated by rhododendron (*Rhododendron ferrugineum*), and a variety of herbaceous plants. On one of these mountains, two non-native species of trees in the Pyrenees have been planted in the 19th century in small patches: (Spruce (*Picea abies*) and larch (*Larix decidua*)).

For our study, we collected a total of 232 faecal samples from western capercaillies in both Norway and France (Supplementary Table S1). In Norway, 164 samples were collected between 2018 and 2019, during autumn (September to November) and spring (April to June). Collections in Norway were carried out in the following counties; Finnmark (6 samples), Innlandet (106 samples), Troms (23 samples) and Trøndelag (56 samples). In France, 68 samples were collected in spring (April to June) between 2016 to 2019, from the following mountain ranges; Jura (18 samples), Vosges (30 samples), and Pyrenees (Ariège and Haute-Garonne) (20 samples) (Fig 1).

**Fig 1:**
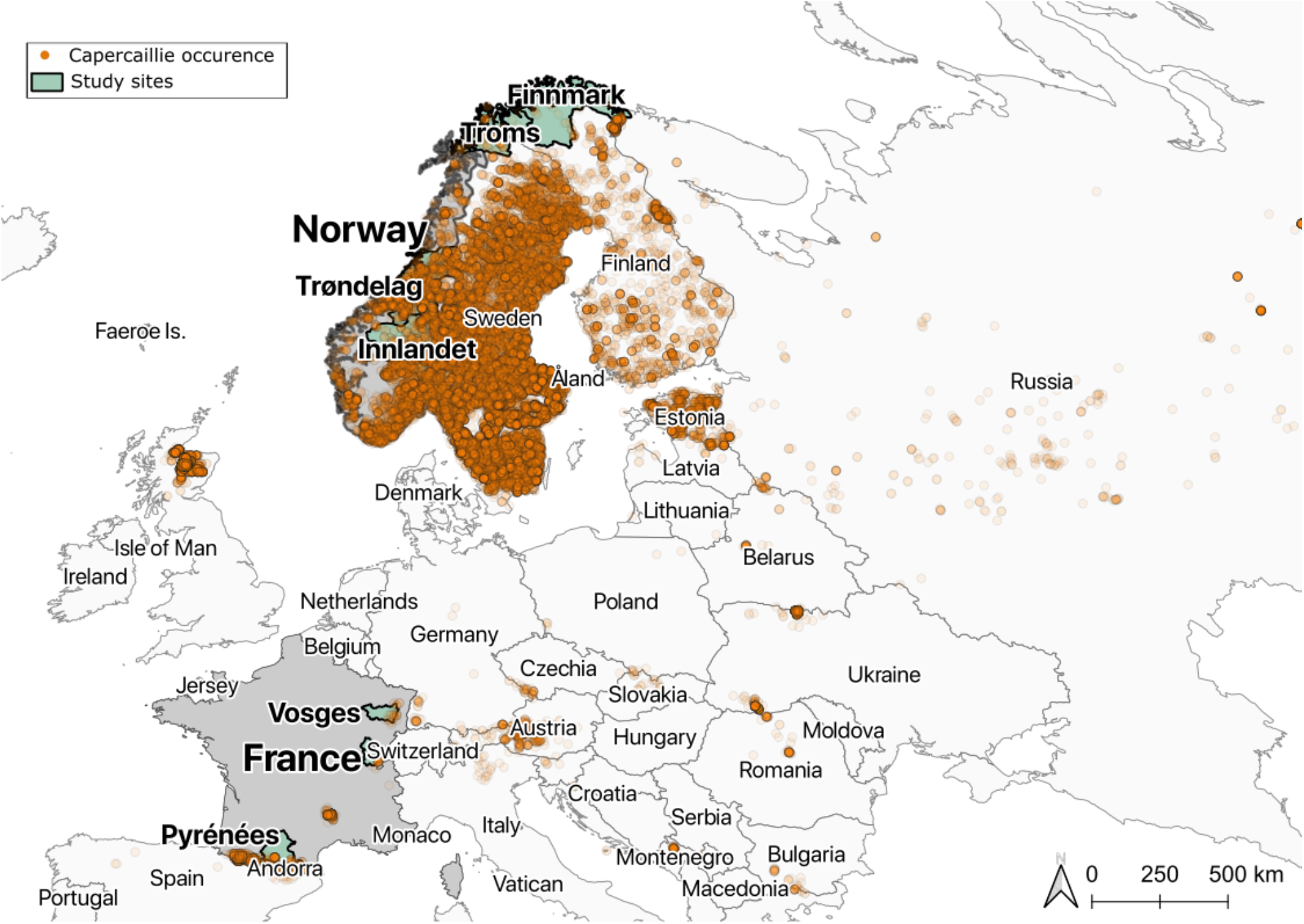
Map showing the occurrence of capercaillies in the past ten years (GBIF.org (04 March 2021) GBIF Occurrence Download https://doi.org/10.15468/dl.ucn5ce), and where capercaillie faecal samples were collected for this study (n=232). Four of the study sites are located in Norway (Finnmark n=6, Troms n=23, Trøndelag n=56, and Innlandet n=106), while three are located in France (Vosges n=30, Jura n=18, and Pyrenees n=20). Created with QGIS version 3.10.4.

Each faecal sample corresponds to one dropping deposited by one individual. Faecal samples were collected in sterile airtight tubes containing Merck silica gel (with colour indicator, granulate size 1-3 mm, Merck KGaA). To minimize the chance of collecting multiple samples from the same individual, only faecal droppings that were minimum 25 m apart were collected (Mollet et al., 2015). All faecal samples were collected opportunistically in the field except one sample, Z01, which was collected from a captive male capercaillie located at the Namsskogan Familiepark (Trøndelag, Norway). For each faecal dropping sample, the sex of the capercaillies who deposited the samples was determined by either visual observation of the defecating individual if present, or by measuring the diameter of the faecal samples (>10 mm for males, <8 mm for females) (Thiel et al., 2008). The faecal samples were stored at −20°C prior to DNA extraction.

### DNA extraction and metabarcoding

DNA extractions were carried out in a dedicated pre-PCR laboratory. For each faecal dropping, approximately 2 mm of the exterior layers of each faeces were removed to reduce environmental contamination. For DNA extraction, approximately 200 mg of faecal material was used for each sample. DNA extractions were carried out following the QIAGEN PowerFaecal DNA Isolation Kit using the manufacturer’s protocol (QIAamp PowerFecal DNA Kit Handbook 08/2017), with the following modification: DNA was eluted in 100 μl Solution C6 with an incubation time of 15 min at 37°C to increase DNA yield. DNA extractions were carried out in batches of 25 to 40 samples with one extraction blank (with no faecal material) included in each batch. DNA extracts were stored at −20°C.

To characterise the diet of the western capercaillie, we used a universal primer set for plants targeting the P6 loop of the *trnL* intron; forward primer *trnL-g* (5′-GGGCAATCCTGAGCCAA-3′) and reverse primer *trnL-h* (5′-CCATTGAGTCTCTGCACCTATC-3′) (marker length: ~10bp to 143bp, excl. primers) (Taberlet et al., 2007). Using *oligotag*, 15 tags of six nucleotides in length were designed and added to the 5′ end of the forward and reverse primer. The tags were designed to have a minimal hamming distance of three. An additional one or two nucleotides were added to the 5′ end of each tag to increase complexity on the flow cell (De Barba et al., 2014). To screen for PCR inhibition and determine the number of cycles for PCR amplification (Murray et al., 2015), we performed quantitative PCR (qPCR) on a dilution series (1:1, 1:5, 1:10) of a subset of the sample DNA extracts (25 samples) and on the positive control (leaf DNA extracts from *Cinchona officinalis*, a species that is not native to Europe). Undiluted extraction and qPCR negative controls were included in the qPCRs. The 20 μl qPCR reactions contained 1 μl of DNA template, 0.75 U AmpliTaq Gold DNA Polymerase, 1x AmpliTaq Gold buffer and 2.5 mM MgCl_2_ (all from Applied Biosystems, USA), 0.2 mM of each dNTP (Invitrogen), 0.5 mg/ml Bovine Serum Albumin (BSA); Bio Labs), 1 μl of SYBR Green/ROX solution (one-part SYBR Green I nucleic acid gel stain (S7563; Invitrogen), four parts ROX Reference Dye (12223-012; Invitrogen), and 2000 parts high-grade DMSO, and 0.4 μM of each of the tagged forward and reverse primers. Quantitative PCR (qPCR) was performed on a MX3005 qPCR machine (Agilent) with the following cycling conditions: 95°C for 10 min, 40 cycles of 95°C for 30 sec, 55°C for 30 sec followed by a dissociation curve (denaturation at 95°C for 1 min, annealing at 55°C for 30 sec and denaturation 95°C for 30 sec). The qPCR screening indicated that using 1 μl of neat sample DNA extract and running 35 cycles were optimal for the subsequent tagged PCR amplification. Amplification and dissociation curves confirmed that only primer-dimers, and not target DNA, were present in the negative extraction controls. Tagged PCRs were carried out with three PCR replicates for each DNA extract, extraction negative control, positive control, and PCR negative control. A total of eight extraction negative controls, 10 PCR negative controls, and seven positive controls (all the same *Cinchona officinalis* DNA extract) were included in the study. PCR replicates were tagged using non-matching nucleotide tag combinations (e.g. forward primer tag 1 - reverse primer tag 2, forward tag 2 - reverse tag 3). Tagged PCRs were carried out as in the qPCR, although omitting the SYBR Green/ROX solution, adding an extension time of 72°C for 7 min instead of the dissociation curve, and using 35 PCR cycles. For PCR blanks, 1 μl of AccuGENE Molecular Biology Water was included instead of template DNA. Up to 2 PCR negative controls were used per PCR reaction. Visualisation of amplified PCR products was done on a 2% agarose gel with GelRed against a 50 bp ladder. Amplification was considered successful if PCR products were between 60 bp and 200 bp when visualised on the agarose gel. Extraction and PCR negative controls did not show any bands on the agarose gel.

Prior to library building, PCR products were kept for pooling if at least two of the three PCR replicates for each sample were successfully amplified. PCR products were pooled together without repeated nucleotide tag combinations. This resulted in six amplicon pools, each containing 72 to 124 PCR replicates with different nucleotide tag combinations within each pool. Extraction, PCR negative controls and positive controls were included in each amplicon pool. We purified the six amplicon pools using MagBio HighPrep beads (2.8× bead to amplicon pool ratio), eluted in 50 μl of EB buffer (Qiagen) and quantified using a Qubit Flex Fluorometer dsDNA HS Assay (Invitrogen). We followed the Tagsteady library preparation protocol to prepare the purified amplicon pools for sequencing (Carøe & Bohmann, 2020). Amplicon libraries were purified using MagBio HighPrep beads (1.2× bead to library ratio) and eluted in 30 μl of EB buffer. Quantification of purified amplicon libraries was carried out with qPCR, using the NEBNext Library Quant Kit for Illumina (New England BioLabs Inc). The six purified amplicon libraries were pooled in equimolar ratios and sequenced using 150 paired-end chemistry on the Illumina Miseq V3 sequencing platform (with 15% PhiX) at the GeoGenetics Sequencing Core, University of Copenhagen, aiming at 35,000 paired reads per PCR replicate.

### Data analysis

Sequence data were analysed using the OBITools package (Boyer et al., 2016). Prior to data processing with OBITools, paired reads were merged using SeqPrep (https://github.com/jstjohn/SeqPrep, v1.2) with default parameters. Sequences were assigned to PCR replicates with *ngsfilter* using a 100% match to tags and maximum one bp mismatch to primers. *Obiuniq* was then used for merging strictly identical sequences. Denoising was carried out with *obigrep*: sequences with only a single copy (singletons) and/or shorter than 10 bp were removed. Amplification and sequencing errors were identified using *obiclean* using a threshold ratio of 5% between the main ‘head’ sequence and the putative erroneous ‘internal’ sequence (Bellemain et al., 2013; De Barba et al., 2014). Sequences identified as ‘internal’ were removed, keeping only ‘head’ sequences. Taxonomic assignment was carried out with *ecotag,* using the global EMBL reference database (release r143 from April 2020) (https://www.ebi.ac.uk/) that was generated using *ecopcr*, and with a local reference database (ArctBorBryo) containing 2280 reference sequences (1053 unique) of the *trnL* p6 loop from 2001 different arctic and boreal vascular plants and bryophytes (Soininen et al., 2015; Sønstebø et al., 2010; Willerslev et al., 2014). For all samples, we prioritised matches against the local ArctBorBryo reference database as it has better accuracy identifying species found in the boreal area (Alsos et al., 2016) but no local reference database was available for France. Post-OBITools sequence filtering and merging of the taxonomic assignments were carried out with a custom R script (https://github.com/Y-Lammers/MergeAndFilter). To balance the removal of false positive and negatives, we used an intermediate stringent filtering in which we keep the sequences fulfilling all of the following criteria: i) matched 98% to either reference database (using 98% rather than 100% allows identification of taxa missing in either reference library), ii) had a minimum of 3 reads for each taxon observed within a PCR replicates, and iii) occurring in at least two PCR replicates of a sample (Alsos et al., 2018; Ficetola et al., 2015). Following this sequence filtering, samples with less than 100 total reads were discarded. Remaining sequences fell into five categories: 1) the taxon identifications from both databases were identical, 2) the taxon identifications matched at the family level but differed at the genus and/or species level, 3) the sequence was identified with only one database and not found in the other, and 4) the taxon was determined to be a contaminant/not plausible based on the known distribution of the taxon. The identified taxa for sequences in category 2 were based on matches to ArctBorBryo if it had the same or higher percentage identity than EMBL. Otherwise, identified taxa were based on matches to EMBL. Sequences in category 4 were removed as false positives. We also checked the sequences found in extraction and PCR negative controls for possible contamination. Sequences that were found only in positive controls or with higher number of reads in negative controls than in samples were removed (Supplementary Table S2). As the P6 loop contains homopolymers, we distinguished *Vaccinium* species based on the homopolymer-A region and merged other taxa according to homopolymers (Supplementary Table S3). Homopolymers are defined as sequences with five or more of the same nucleotide. The resulting list of plant taxa was grouped according to the following eight functional groups: trees, shrubs, dwarf shrubs, forbs, graminoids, aquatics, vascular cryptogams or bryophytes. Additionally, we checked that the assigned taxa were known from the geographical region using reference material online (https://www.biodiversity.no/, https://www.gbif.org/).

Statistical analyses were carried out using R (version 3.6.2). Diet richness analyses were based on the presence/absence of each plant taxa in each faecal sample and calculated as frequency of occurrence (F_O_%) (Quéméré et al., 2013). For analysis of compositional differences, we used relative read abundance (RRA), which is the proportion of identified reads assigned to each plant taxon (Kartzinel et al., 2015; Soininen et al., 2009; Willerslev et al., 2014). Read counts were first transformed into RRA data using the *vegan* package with *decostand* function (Oksanen et al., 2015). Dissimilarities in dietary composition between faecal samples collected from three variables (locations, seasons, and sex) were quantified using the Bray-Curtis distance, with K = 3 dimensions and 999 maximum iterations through the *metaMDS* function in *vegan (Bray & Curtis, 1957*). Statistical differences in dietary compositional dissimilarities were computed using the permutation multivariate analysis of variance (perMANOVA) (999 permutations), with the *adonis* function (Anderson, 2001). The *betadisper* and *permutest* function with 999 permutations and bias adjustment was used to test for the perMANOVA assumption of homogeneity of multivariate dispersion (intraspecific dietary variation). For visualization of patterns, we used the nonmetric multidimensional scaling (NMDS) plot (Shepard, 1962a, 1962b) with an acceptable stress level of <0.2 (Clarke, 1993). We also used the *envfit* function with 999 permutations to investigate which plant taxa may be involved in driving distribution patterns. Additionally, we ran an indicator taxa analysis using the *indicspecies* package to determine which plant taxa were strongly associated with a given variable (De Cáceres & Legendre, 2009). To determine the number of potential new diet items, duplicated taxa were merged according to taxon name (i.e Pinus1, Pinus2), resulting in 95 unique plant taxa (Supplementary Table S4).

## RESULTS

### Final dataset description

After removing 60 samples dataset due to either unsuccessful PCR amplification (53 samples, 20 of them from one batch of samples collected from Innlandet, Norway (V01 - V34)) or filtering parameters (7 samples) used in data analysis (Supplementary Table S5), our final dataset consisted of 172 western capercaillie faecal samples. From these, 163 sequences with a minimum percentage identity of 98% to at least one taxon in the ArctBorBryo and/or EMBL reference database (16,477,6339 reads) were kept after taxonomic assignment. Of the 163 sequences, 95 sequences had 100% match to ArctBorBryo (1,5010,048 reads), 136 sequences had 100% match to EMBL (16,386,507 reads), and 92 sequences had 100% match to both reference database (15,003,823 reads). The remaining sequences matched at 98% to either database and the non-matching part of the sequences primarily consisted of homopolymers (84,607 reads). Of the 163 sequences, 34 (20.86%) were assigned to family level, 59 (36.20%) to genus, and 67 (41.10%) to species level. The remaining three sequences (1.84%) were assigned to higher taxonomic levels (2 orders and 1 subclass). These three sequences (sequence ID: 4667, 6932, and 2048) were kept as they belonged to the bryophytes functional group, and bryophytes sequences are less well represented in *trnL* P6 loop reference databases.

### Dietary richness

After merging of homopolymers, a total of 122 plant taxa were identified, with 119 taxa identified to at least family level; these comprised 46 families, 68 genera, and 46 species from eight functional groups (trees, shrubs, dwarf shrubs, forbs, graminoids, aquatics, vascular cryptogams or bryophytes; Table 1). Of the taxa, 37.7% could be identified to species level, 33.6% at the genus level, and 28.7% at the family level. Based on the frequency of occurrence among samples, the most frequently occurring groups of plants were trees (39.06%), followed by forbs (16.41%) and dwarf shrubs (16.11%). The least occurring plants were aquatics (0.15%). Shrubs (5.02%), bryophytes (6.84%), vascular cryptogams (7.6%), and graminoids (8.81%) made up the rest of the plants found in capercaillie faecal samples (Supplementary Fig S1.1). The two most frequently occurring plant families consumed were Pinaceae (97.69% occurrence across all samples) and Ericaceae (71.1%) (Table 1). The most common taxon of the Pinaceae family was Pinus (58.78% of all Pinaceae), while *Vaccinium myrtillus* (54.72%) was the most commonly identified taxon from the Ericaceae family (Supplementary Fig S1.2). Between autumn and spring, there was a significant difference in the F_O_ % of both Pinaceae and Ericaceae found across all samples (Two sample T-test, p-value <0.05).

**Table 1:** Frequency of occurrence (F_O_) of plant families found across all samples (n = 172)

| Functional groups | Family | No. of occurrences<br>in samples | Frequency of<br>occurrence (F<sub>o</sub> %) |
| --- | --- | --- | --- |
| Trees | Betulaceae | 86 | 50.0 |
|  | Fagaceae | 39 | 22.7 |
|  | Malvaceae | 4 | 2.3 |
|  | Pinaceae | 168 | 97.7 |
|  | Salicaceae | 27 | 15.7 |
| Shrubs | Cupressaceae | 62 | 36.0 |
|  | Hypericaceae | 3 | 1.7 |
|  | Juncaceae | 11 | 6.4 |
| Dwarf Shrubs | Ericaceae | 122 | 70.9 |
| Forbs | Apiaceae | 6 | 3.5 |
|  | Asteraceae | 15 | 8.7 |
|  | Brassicaceae | 11 | 6.4 |
|  | Campanulaceae | 2 | 1.2 |
|  | Caryophyllaceae | 6 | 3.5 |
|  | Chenopodiaceae | 1 | 0.6 |
|  | Convolvulaceae | 1 | 0.6 |
|  | Cornaceae | 4 | 2.3 |
|  | Fabaceae | 10 | 5.8 |
|  | Geraniaceae | 7 | 4.1 |
|  | Lamiaceae | 1 | 0.6 |
|  | Nartheciaceae | 1 | 0.6 |
|  | Orobanchaceae | 9 | 5.2 |
|  | Oxalidaceae | 11 | 6.4 |
|  | Plantaginaceae | 2 | 1.2 |
|  | Polygonaceae | 5 | 2.9 |
|  | Ranunculaceae | 31 | 18.0 |
|  | Rosaceae | 41 | 23.8 |
|  | Rubiaceae | 1 | 0.6 |
|  | Saxifragaceae | 2 | 1.2 |
| Graminoids | Cyperaceae | 21 | 12.2 |
|  | Poaceae | 38 | 22.1 |
| Aquatics | Menyanthaceae | 2 | 1.2 |
| Vascular cryptogams | Athyriaceae | 15 | 8.7 |
|  | Blechnaceae | 10 | 5.8 |
|  | Cystopteridaceae | 24 | 14.0 |
|  | Dryopteridaceae | 11 | 6.4 |
|  | Equisetaceae | 18 | 10.5 |
|  | Lycopodiaceae | 13 | 7.6 |
|  | Thelypteridaceae | 6 | 3.5 |
| Bryophytes | Dicranaceae | 19 | 11.0 |
|  | Hylocomiaceae | 4 | 2.3 |
|  | Hypnaceae | 3 | 1.7 |
|  | Mniaceae | 4 | 2.3 |
|  | Polytrichaceae | 12 | 7.0 |
|  | Sphagnaceae | 25 | 14.5 |
|  | Splachnaceae | 2 | 1.2 |

From each faecal sample, we identified on average seven plant taxa (SD:+− 5, range: 1–27). Only 2.33% of samples contained just one plant taxa (monodiets). The average dietary richness for samples collected between locations varied from 5.7 taxa in Troms, to 9.7 taxa in Finnmark (Table 2). There were no significant differences in dietary richness between samples collected from the seven different locations (One-way ANOVA, p-value = 0.0716), between sexes (Two sample T-test, p-value = 0.958) nor between seasons (Two sample T-test, p-value = 0.068). Of the 122 plant taxa retrieved, capercaillies located in Trøndelag fed on the largest diversity of taxa identified at 87 taxa, while the lowest diversity of plants identified were from the Jura population at 23 taxa (Table 2). Additionally, we retrieved 23 plant taxa that were found only in female capercaillie’s faecal samples, while 16 plant taxa were found only in the male capercaillie’s faecal samples (Supplementary Table S6). Season-wise, 27 plant taxa were identified only from spring samples, while 25 plant taxa were identified only from autumn samples.

**Table 2:**
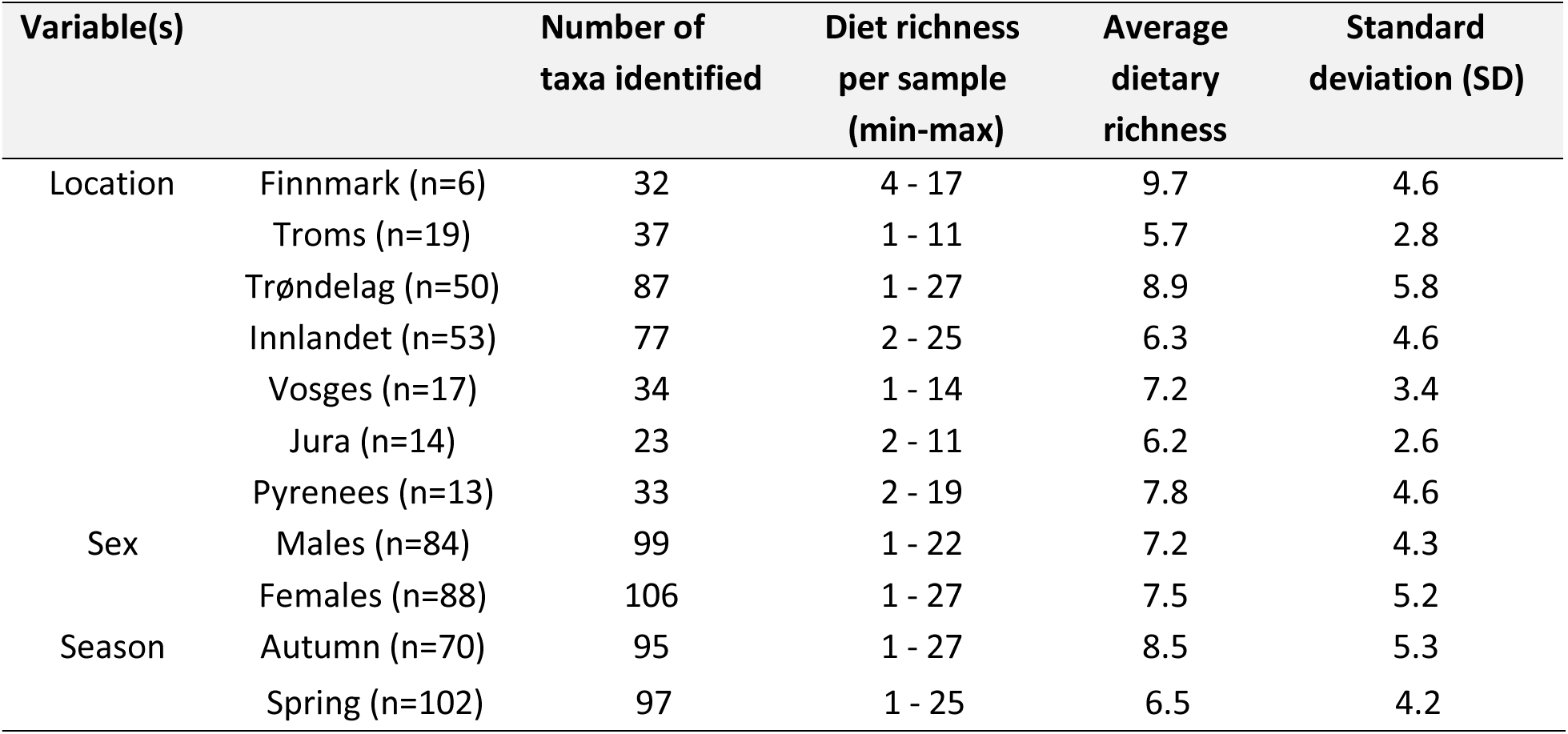
Dietary richness obtained from capercaillie faecal samples collected for each variable, with the number of taxa identified, minimum (min) and maximum (max) diet richness range, average, and standard deviation.

### Dietary composition

Similar to the dietary richness data, the highest relative read abundance (RRA) of consumed plant taxa were trees, dwarf shrubs, and forbs, making up 66.47%, 15.76%, and 7.50% of the total RRA, respectively (Supplementary Fig S2.1). Plant taxa belonging to these three functional groups were found in samples collected in all locations, seasons, and in both sexes (Supplementary Fig S2.2 and S2.3). The least consumed plants were aquatics (RRA 0.02%). The two families with the highest RRA were Pinaceae (56.85%) and Ericaceae (15.76%), and the main taxa making up the Pinaceae family was Pinus (70.25% of all Pinaceae), while *Vaccinium myrtillus* (65.44%) formed the majority of Ericaceae (Supplementary Fig S2.4). There was a significantly higher proportion of RRA of Pinaceae found in samples from spring (average 0.42%) than autumn (average 0.21%, Two sample T-test, p-value <0.05), but there was no significant difference for Ericaceae (Two sample T-test, *p*-value = 0.39) (Supplementary Fig S2.5). The proportion of reads assigned to the plant functional groups at each location, separated by sex and season, is visualised in Fig 2.

**Fig 2:**
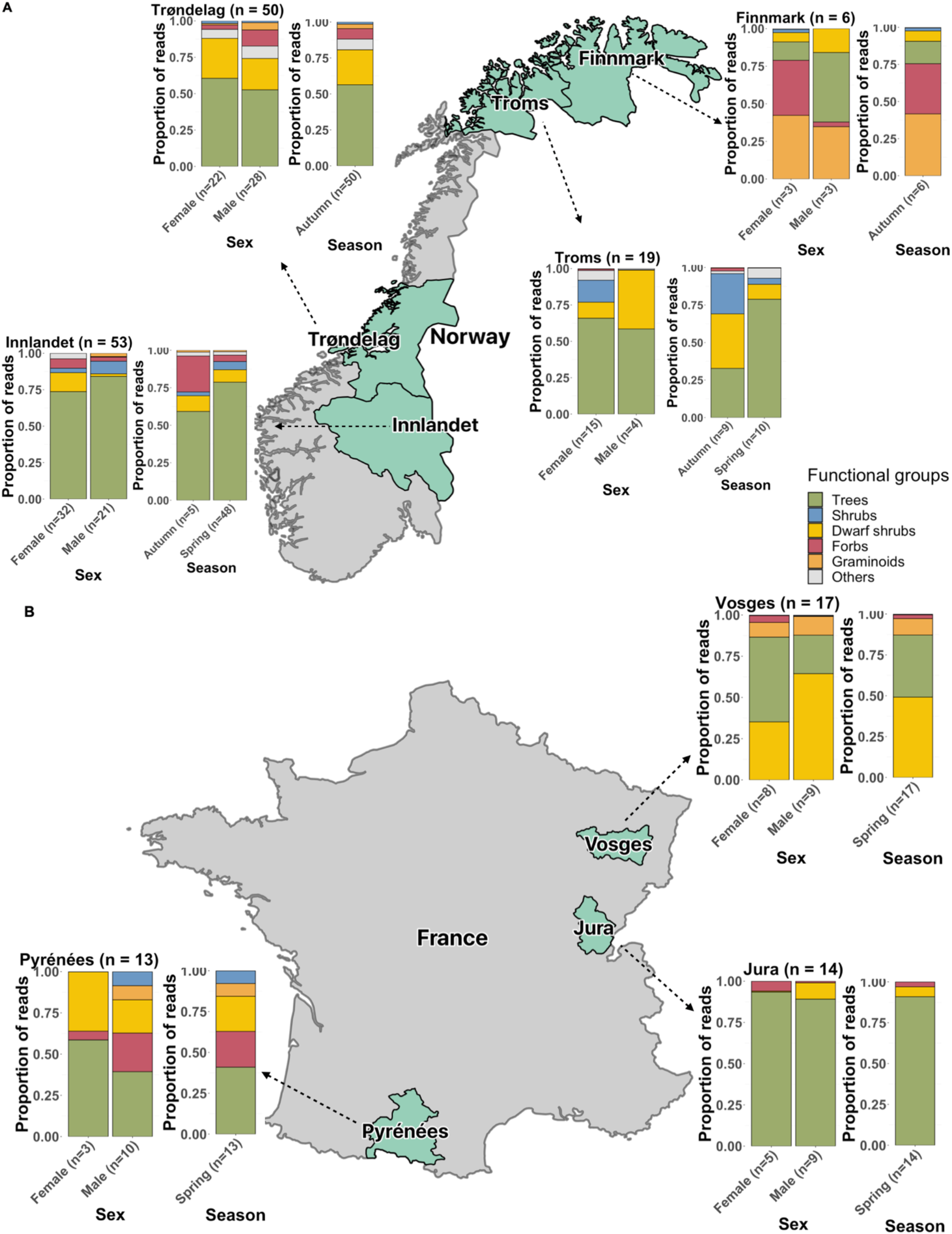
Stacked bar charts showing the proportion of reads assigned to functional groups between each location, sex, and season from capercaillie faecal samples collected in **a)** Norway, and **b)** France.

The perMANOVA test (with Bonferroni *p*-value correction) for the effects of variables (location, sex, and season) on the dietary composition based on RRA at the level of plant taxa showed that there was a significant difference between faecal samples collected from the different locations (adonis F_5,,86_ = 11.01, *r^2^* = 0.17, *p* = 0.001) and between seasons (adonis F_2,03_ = 0.64, *r^2^*= 0.01, *p* = 0.036) (Fig 3a). There were no significant differences in dietary composition between males and females across all faecal samples (adonis F_1,63_ = 0.51, *r^2^* = 0.01, *p* = 0.095). However, there was a significant difference in dietary composition between faecal samples collected from males and females at each individual location (adonis F_1,47_ = 2.77, *r^2^* = 0.04, *p* = 0.024) (Fig 3b).

**Fig 3:**
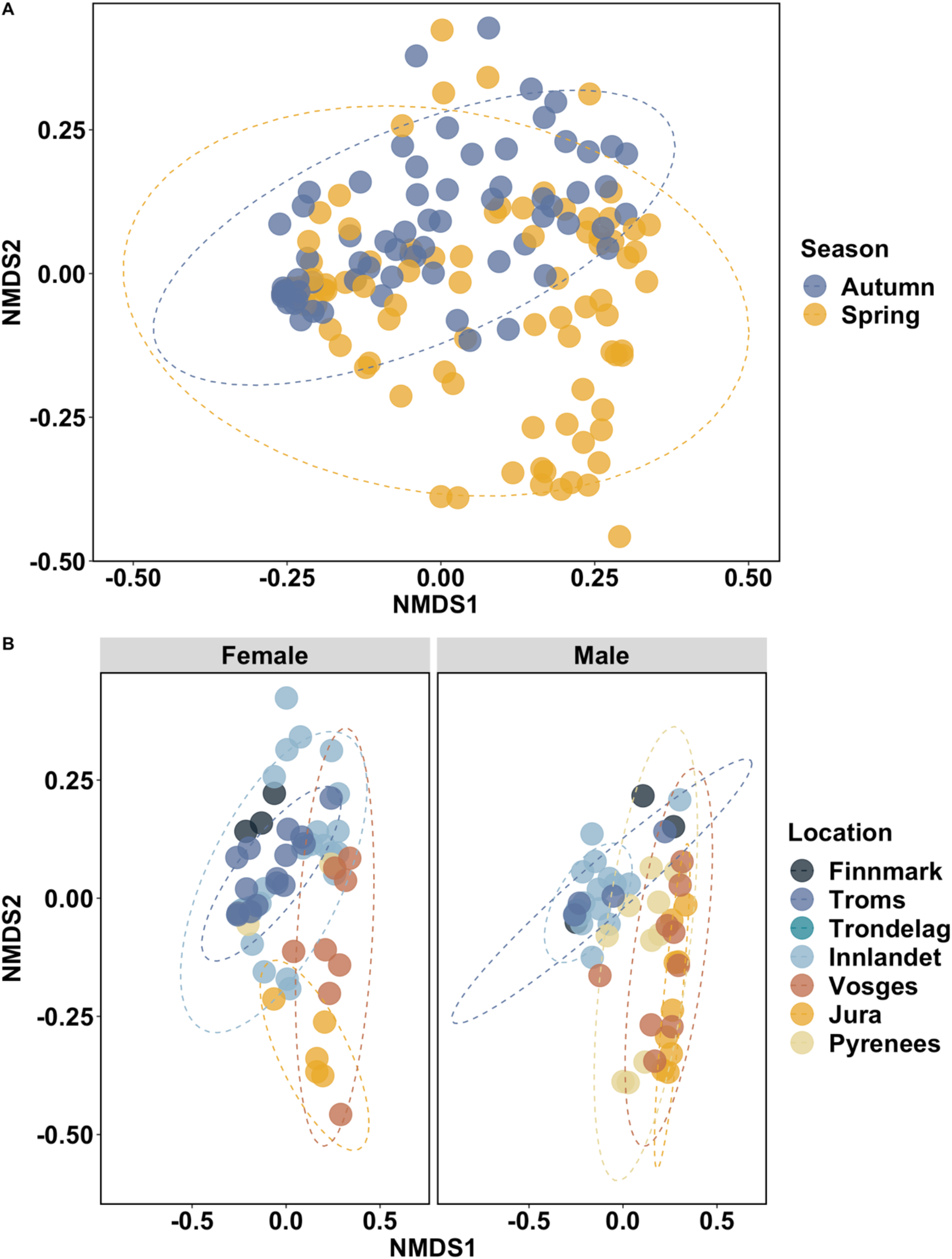
NMDS plots of RRA-based Bray-Curtis dissimilarity of capercaillie faecal samples collected from **a)** different seasons (adonis F_2.03_, = 0.64, *r^2^* = 0.01, *p* = 0.036), and **b)** both sexes from each of the seven locations (adonis F_1,47_, = 2.77, *r^2^* = 0.04, *p* = 0.024). The stress level of 0.163 is under the cut-off value of 0.2 for an interpretable ordination as suggested by Clark (1993).

The test for perMANOVA assumption of homogeneity of multivariate dispersion returned insignificant pseudo F and permutative p-values > 0.05, showing that while there is a difference in composition, there was no dispersion between variables tested which supports the assumption of homogeneity. The *envfit* test to uncover which plant taxa drive these results showed 16 plant taxa that were statistically significant (*p* ≤ 0.05) (Supplementary Table S7, Fig S2.6). From the indicator taxa analysis based on locations, 20 plant taxa were strongly associated with a single location, while one taxon was strongly associated with four locations (Troms, Trøndelag, Innlandet, and Pyrenees) (*p* ≤ 0.05, Supplementary Table S8). Two plant taxa were strongly associated with samples collected from females, while three plant taxa were strongly associated with samples collected from males. For the seasons, 19 plant taxa were strongly associated with samples collected from autumn, while two plant taxa were strongly associated with samples collected from spring (Supplementary Table S8).

## DISCUSSION

Dietary information is a critical component in understanding animal ecology and effective conservation management. Through metabarcoding of 172 capercaillie faecal samples, we reconstructed a list of plant taxa (n= 122) consumed by capercaillies. All except three taxa were identified to minimum family level, and more than one-third of the taxa were identified at the species level (37.7%). We highlighted the differences in dietary composition of populations located across the seven locations, between seasons and between sexes at each of the seven locations. Additionally, we also identified new diet items that were previously not recorded based on non-molecular methods, showing the potential of metabarcoding in retrieving a wider range of diet items, and at a better taxonomic resolution. From this, we learnt that capercaillie feeds on a wider range of plants than expected, suggesting that they can adapt their diet to resource availability. This ecological information can be incorporated into the planning of appropriate conservation strategies for the western capercaillies.

Differences in diet observed between the analysed capercaillie populations and seasons are expected, given that the vegetation varies from location and are subject to seasonal effects (Jacob, 1987; Saniga, 1998). Moreover, diet plasticity and individual preferences have been documented even within a small geographical location, and only few plant taxa have been recognised as being important diet items throughout their range (Gonzalez et al., 2012; Sedinger, 1997). The most frequently occurring group of plants with the highest RRA found in the analysed capercaillie faecal samples were trees, dwarf shrubs, and forbs. Pinaceae and Ericaceae were the most frequently occurring families. This result is in accordance with previous studies on capercaillie diet based on non-molecular methods, which have shown that plants from these two plant families form the main components of their habitats and diet throughout the season, but its importance shifts with changes in the season (N. Picozzi et al., 1996; Ilse Storch, 1993). During seasons where snow cover is greater such as in winter or early spring (Gjerde & Wegge, 1989), pine needles are a critical resource for survivability when ground vegetation is scarce. By contrast, during summer and autumn, dependence on pine needles decreases as the availability of dwarf shrubs from the Ericaceae species, specifically bilberries (also called European blueberries, *Vaccinium myrtillus*), increases. This shift in diet consumption - from pine needles in the cold seasons to blueberries in the warmer seasons - has been well documented in non-molecular studies and are important food items across their range (Blanco-Fontao et al., 2010; Quevedo et al., 2006; Rodríguez & Obeso, 2000; Ilse Storch, 1993). Also, in the present study, we show Pinaceae consumption is highest in spring when snow often still covers the ground, while Ericaceae consumption shows the opposite pattern with highest prevalence in the diet during autumn.

We had expected autumn samples to have a higher diet variety than spring samples, due to the presence of snow cover in spring which reduces access to ground vegetation. However, samples collected during the two seasons had almost the same number of taxa identified. This could be due to differences in snow cover between the individual locations even during the same collection period, making comparisons between seasons at different sites difficult. The almost complete absence of bryophytes and vascular cryptogams in the diet of the French capercaillie populations was also unexpected, given that mosses and ferns have been documented as part of their diet (Blanco-Fontao et al., 2010; Jacob, 1987; Rodríguez & Obeso, 2000), and are present even in early spring. This may at least be partially explained by the strong presence of the Fagaceae trees (beech) which start to bud in winter and were found in 66.7% of the French samples. Studies have shown monodiet feeding on beech buds for several months in spring, as it is a more convenient source of food than mosses and ferns where snow cover is still heavy (Blanco-Fontao et al., 2010; Rodríguez & Obeso, 2000). Another reason could be that the *trnL* P6 loop reference data for bryophytes are not well represented, hence are not identified.

Monodiets, where only a single food item is consumed, is considered a common foraging pattern of capercaillies. During such foraging, they will perch for days or even weeks on individual trees to feed (Sedinger, 1997). In our study, we observe such a monodiet in only 2.33% of the faecal samples collected, with faecal samples containing on average eight different plant taxa. This may signal that the western capercaillies are moving more frequently over longer distances while foraging, at the expense of increasing energy expenditure to energy-intake (Blanco-Fontao et al., 2010). Alternatively, metabarcoding may be a more sensitive and powerful method to detect a larger variety of diet items within one faecal sample. This is supported by studies that have directly compared non-molecular methods and metabarcoding for reconstructing diet from the same individuals (Castle et al., 2020; Nichols et al., 2016).

Overall, there were no significant differences in the diet between males and females. However, the diet of females was strongly associated with *Alnus* (from the Betulaceae family) which corresponds to a habitat partitioning study showing a strong association between females and birch trees (Betulaceae) (Bañuelos et al., 2008). Moreover, when comparing the sexes within each location, there was a significant difference in the dietary composition between males and females. As grouses are a sexually dimorphic family that exhibits habitat partitioning throughout the seasons (Connelly, 1989), we had expected to see this difference given that capercaillies are the most sexually dimorphic of all grouses (Bañuelos et al., 2008) and female reproductive needs would result in differential habitat and resource preferences as compared to males (Ilse Storch, 1993).

Due to the differences in diet observed throughout the capercaillies range, there is a need to collect localised data to develop effective conservation strategies. Important diet items may need to be protected in the capercaillie habitats, e.g. by controlling grazing from livestock to aid in population recovery and prevent competition for food which could lead to an increase in foraging time and hence, greater exposure to predators (Selås, 2000). Particularly in France (Vosges and Jura), where the capercaillie populations have reduced significantly in the past few decades despite ongoing conservation efforts (Montadert, 2012; Sachot et al., 2006; Segelbacher et al., 2003), our study provides insights into resource use. Here, conservation measures may be tailored to include our findings, by including a focus on preserving the most important diet items for this species. Nevertheless, the wide range of diet items retrieved from our study also indicates that vegetation may not limit food intake for capercaillies, suggesting dietary plasticity in this species. Such plasticity suggests that threats to their survivability may not be resource-related, but rather, other factors such as chick predation or disturbances at mating grounds and during breeding season may play a more pertinent role in the species’ decline (Brenot et al., 1996; Wegge & Kastdalen, 2007).

Utilising metabarcoding for the reconstruction of capercaillie diet has allowed us to retrieve more diet items (122 taxa) and at a higher taxonomic resolution based on 172 samples than non-molecular techniques. Of these 122 taxa, 35 are potentially new diet items that have not been documented in decades of capercaillie diet research using non-molecular means. The inclusion of a captive capercaillie in our dataset did not yield any new diet items that were not also consumed by wild populations. To date, non-molecular capercaillie diet studies identified on average around 10-30 taxa per study, even with large sample sizes (Blanco-Fontao et al., 2010; Borchtchevski, 2009; Gonzalez et al., 2012; Jacob, 1987; Odden et al., 2003; N. Picozzi et al., 1996; Nicholas Picozzi et al., 1999; Rodríguez & Obeso, 2000; Rolstad, 1988; Wegge & Kastdalen, 2008). Current dietary information on the western capercaillies is primarily based on non-molecular techniques, and typically limited to a restricted geographic region and/or very small sample sizes. In Norway and France - the focus areas of the present study - capercaillie diet studies have been restricted to chicks or populations located in South-East Norway (Odden et al., 2003; Rolstad, 1988; Wegge & Kastdalen, 2008), or are based on only a few individuals in France (Brezard, 1980; Jacob, 1987). With metabarcoding, the generation of data can come at a much faster pace (Nichols et al., 2016), and also provides a fuller overview of the diet consumed since fully digested plant fragments can be identified. In comparison, non-molecular methods such as microhistology are less sensitive, require trained experts to accurately identify plant fragments and digested fragments are easily missed (Nichols et al., 2016; Sheppard & Harwood, 2005; Shrestha & Wegge, 2006). Moreover, the study by Nichols et al. (2016) estimated that metabarcoding only costs €115 per sample as compared to €300 using microscopy. Thus, non-molecular methods are also much more expensive with the high time-to-salary costs incurred for analysis.

Our study showcases the high potential of metabarcoding in providing more in-depth information for the understanding of species ecology. However, our study also exemplifies that there are some limitations of this method in diet analysis. First, around 22% (53 samples) of samples failed to amplify during the PCR amplification process. We suspect that the main reason for PCR failure was the way that the samples were stored and transported before DNA extraction. Most of the faecal samples with failed PCR amplification were collected fresh after defecation but were not kept cool immediately following collection. Together with fluctuations in temperature during transportation, this may have led to DNA degradation, affecting the yield of recoverable DNA and leading to failed PCR amplification (Nsubuga et al., 2004). Delays incurred during the transportation process could also have exacerbated the DNA degradation process. When working with faecal samples that are rare and difficult to collect, care has to be taken to minimise the risk of DNA degradation, for example by collecting samples in silica gel. Second, another potential issue with metabarcoding is the high sensitivity of the technique, which is great for enhanced taxa detectability as compared to non-molecular methods, but at the risk of artificially inflating dietary breadth due to potential contamination. Contaminants can be introduced at many stages including sample collection, DNA extraction, and PCR amplification (Lusk, 2014; Valentini, Miquel, et al., 2009; Van Geel et al., 2014). The inclusion of extraction and PCR negative controls allowed us to check for possible contaminants and remove them from our dataset. Manual curation of taxonomically identified sequences was also an essential step in removing food contaminants not found in study areas. For example, we removed sequences corresponding to ginger as ginger plants are non-native and were not present at our study sites. Third, the comprehensiveness of the reference database used for taxonomic assignment is an important consideration. The species resolution of taxa detected from the Norwegian population was 37.72% as compared to 27.12% for the French population. This better taxonomic resolution for the Norwegian samples can be explained by the local reference database used for arctic plants, something that is missing for Central-Southern European plants. Hence, we can argue that the power and sensitivity of metabarcoding over non-molecular techniques rely on the reference database available. Last, metabarcoding is unable to discern which part of the plants are consumed, which may be an important aspect of studies determining energy intake and expenditure. Such studies would have to rely on non-molecular means in addition to utilising metabarcoding.

In conclusion, our results provide the most complete view on capercaillie plant diet to date, and highlight the need for dietary studies to be conducted at different spatial and temporal scales in order to obtain a complete view of a species’ diet. The advantages of metabarcoding over non-molecular techniques in studying animal diet supports its use for future diet studies, but careful consideration of its limitations should be taken into account. Depending on research needs, metabarcoding may have to be used in conjunction with other non-molecular techniques in diet studies.

## Author contributions

PYSC, KB, SB, and IGA designed the research; PYSC, TE, and EM collected samples; PYSC performed laboratory work; YL did the bioinformatic analysis; PYSC did the post-OBITools data analysis; PYSC did the statistical analyses; EM provided details on known/unknown diet of capercaillies; PYSC made the figures and wrote the paper with input from YL, TE, EM, KB, SB, and IGA. All authors contributed to the final version of the submitted manuscript.

## Acknowledgements

For generating the Illumina data, we would like to thank the staff at the Danish National High-Throughput Sequencing Centre. We would like to extend our gratitude to Alex Crampton-Platt (NatureMetrics), Arne Flor, Bjørn Morten Baardvik, Joy Coppes, Kat Bruce (NatureMetrics), Marc Montadert (France National agency for wildlife), Stein Nilsen and Stephanie Witczak for providing relevant contact information, details about capercaillies, or contributing to the development of this work. For sample collection, we would like to thank the following group of people without whom this research would not have been possible; Alexandra Depraz (Groupe Tétras Jura), Bjørn-Roar Hagen, Camilla Hjorth Scharff-Olsen, Christophe Lhez, Hans Christian Pedersen, Kevin Foulché, Kristine Vesterdorf, Maria Ariza Salazar, Per Gustav Thingstad, Pål Fossland Moa, Stéphane Roche, Stian Rembjør Almstad, Torbjørn Alm, and Unni Bjerke Gamst. Additionally, we are grateful to Per Wegge and Françoise Preiss (Groupe Tétras Vosges) for invaluable inputs to the writing of this manuscript and for sample collection. This research is part of the H2020 MSCA-ITN-ETN Plant.ID network, and has received funding from the European Union’s Horizon 2020 research and innovation programme under grant agreement No 765000.

## Supplementary information

Supp1 – Supplementary Tables S1 to S12

Supp2 – Supplementary Figures

## Data accessibility statement

All sequenced data and scripts used for generating data are available on the University of Copenhagen Electronic Research Data Archive (UCPH ERDA) Digital Repository, DOI: xxxxxx upon acceptance of the manuscript

