## Supplementary Fig for "Molecular dietary analyses of western capercaillies (*Tetrao urogallus*) reveal a diverse diet"

Supplementary Information (2/2)

**Table of Contents:**

| **S1. Dietary richness** | Page 2 |
| --- | --- |
| **S2. Dietary composition** | Page 4 |
| **References** | Page 9 |

**S1. Dietary richness**

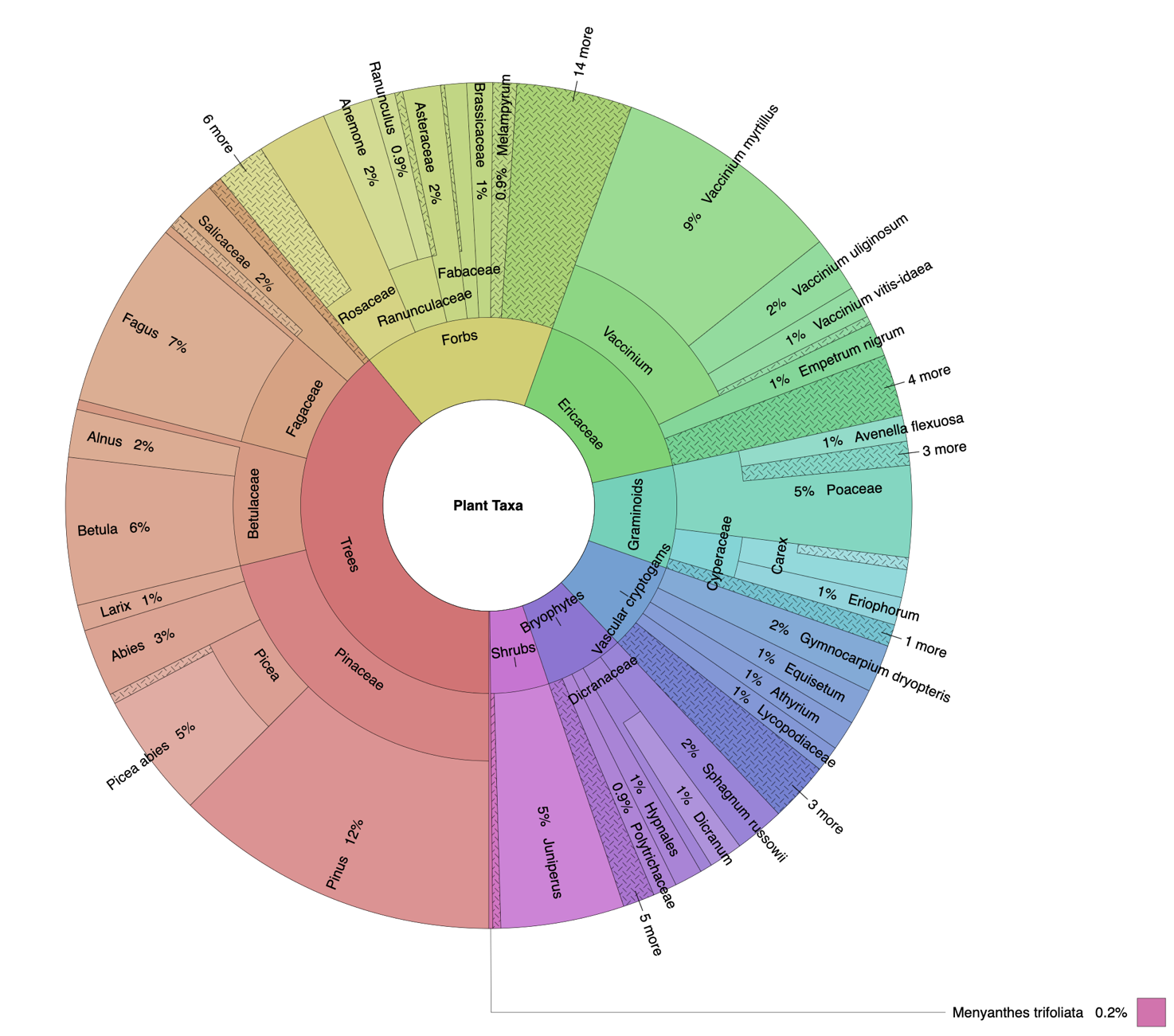

**Fig S1.1:** Krona chart showing the frequency of occurrences of plant taxa retrieved from capercaillie faecal samples collected from Norway and France (n = 172) (Ondov, Bergman, & Phillippy, 2011).

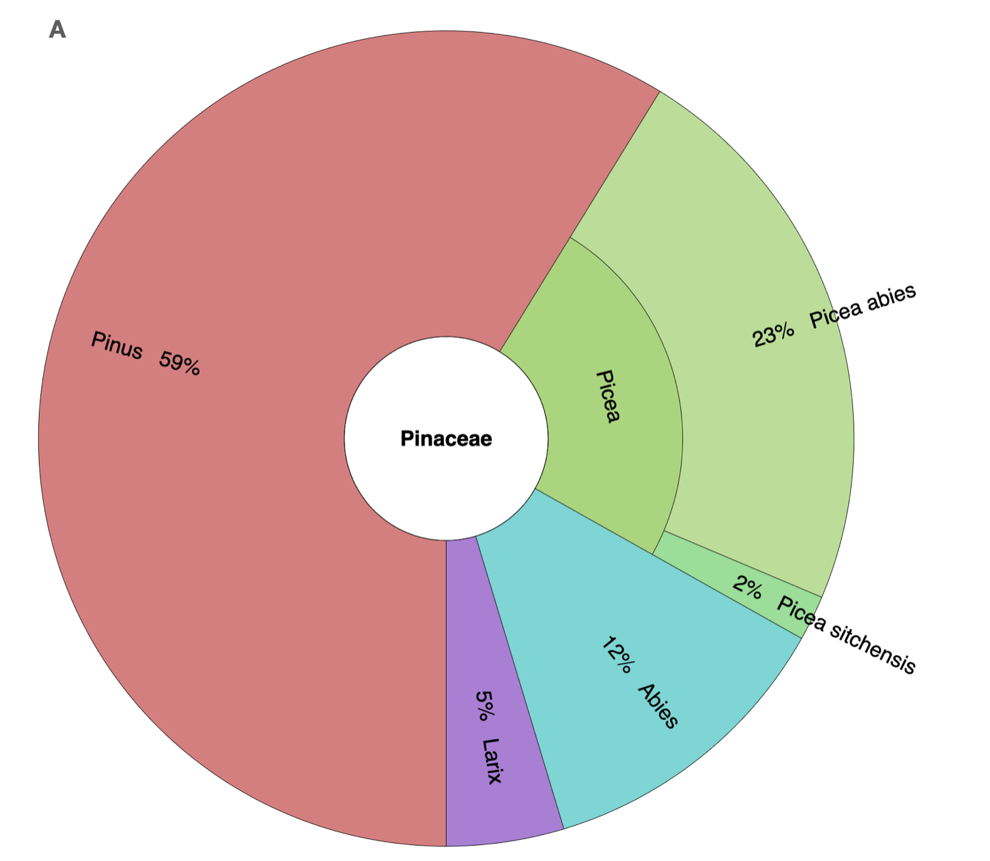

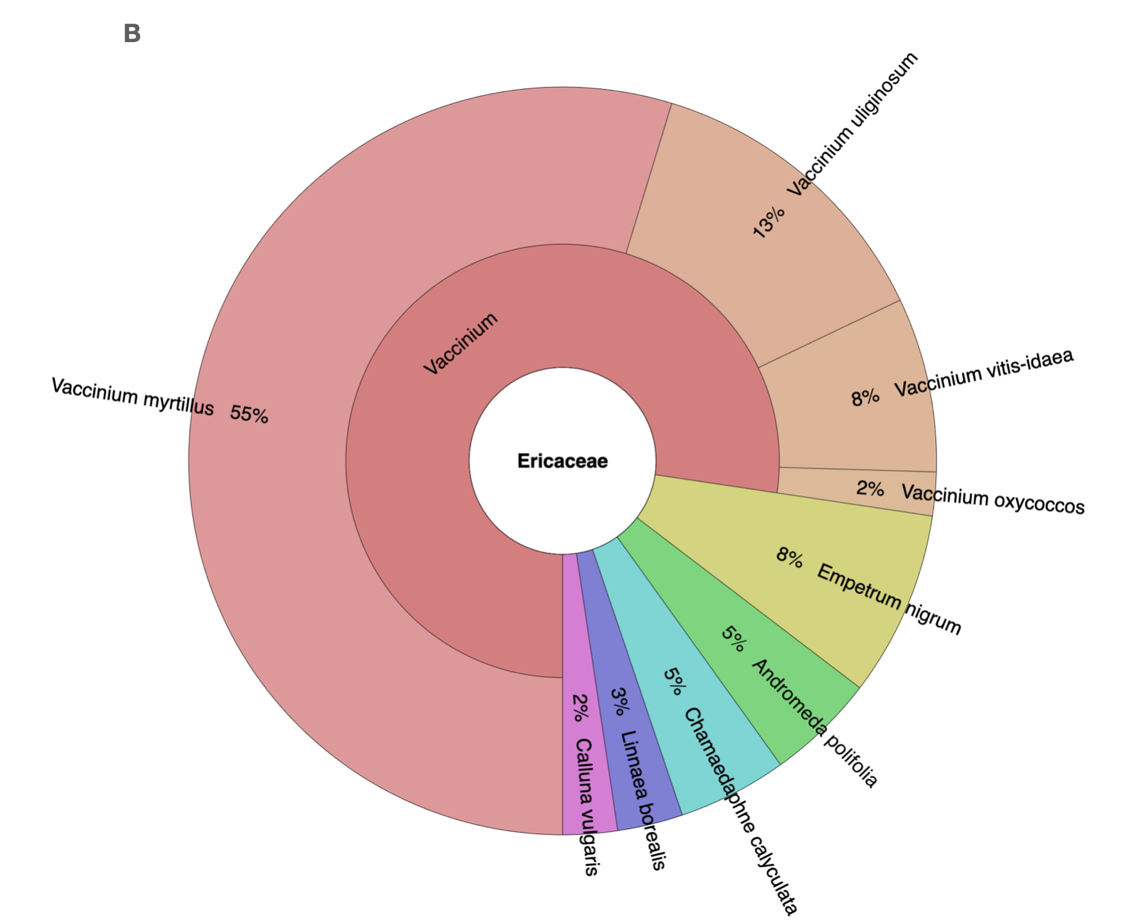

**Fig S1.2:** Krona charts showing the frequency of occurrence of plant taxa found in the families **a)** Pinaceae and, **b)** Ericaceae (Ondov et al., 2011).

**S2. Dietary composition (RRA data)**

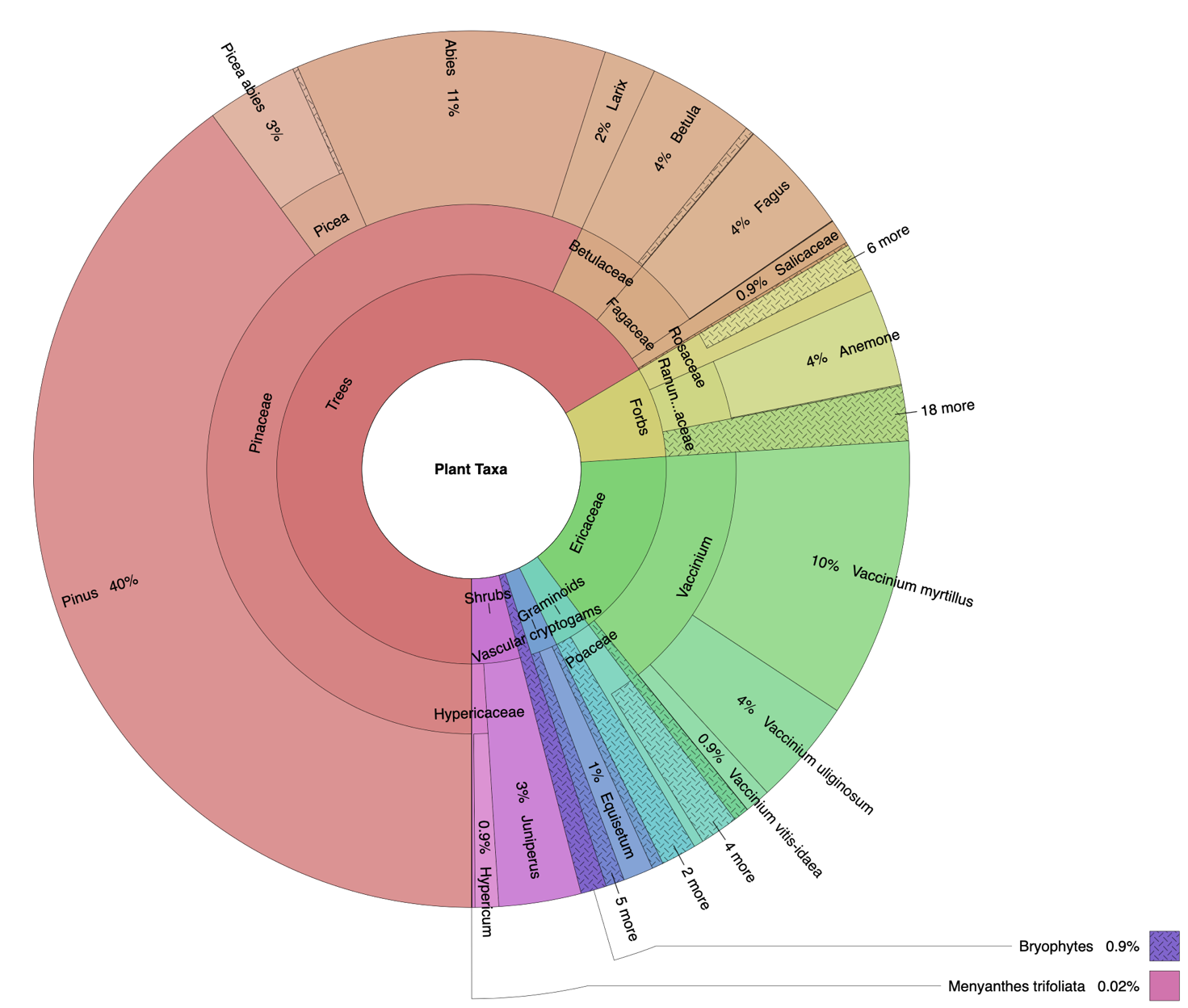

**Fig S2.1:** Krona chart showing the composition of plant taxa based on relative-read abundance (RRA) retrieved from capercaillie faecal samples collected from Norway and France (n = 172) (Ondov et al., 2011).

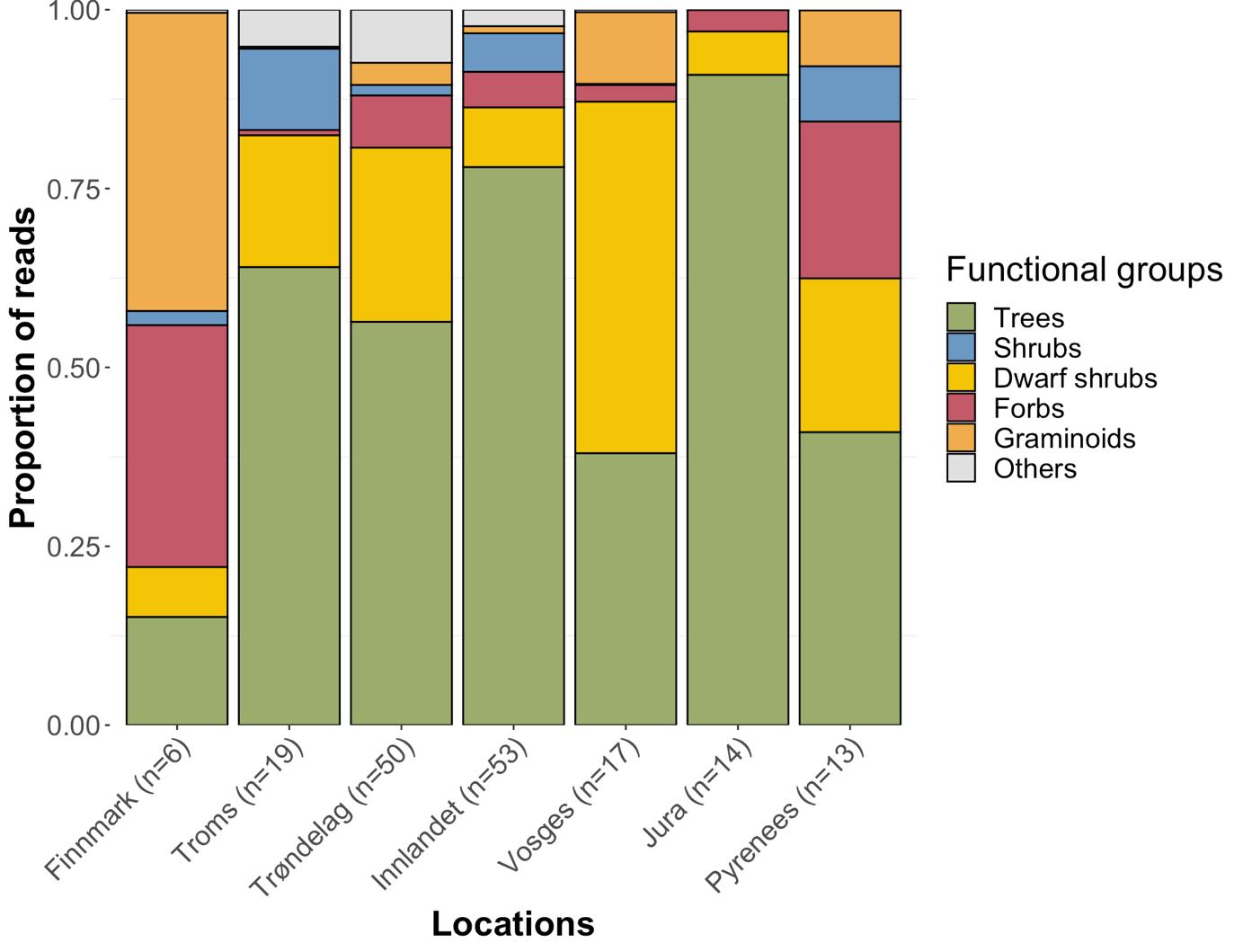

**Fig S2.2:** Dietary composition of plant functional groups retrieved from capercaillie faecal samples collected from Finnmark, Troms, Trøndelag, Innlandet, Vosges, Jura, and Pyrenees.

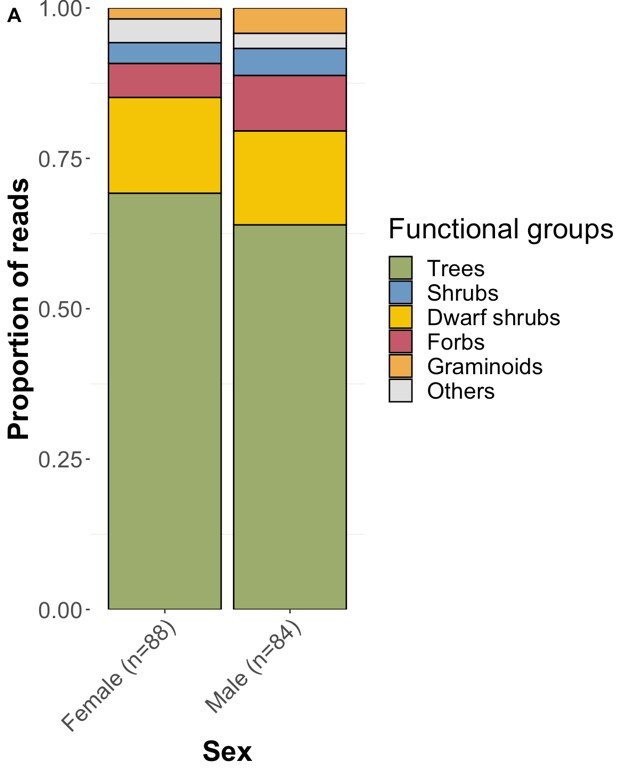

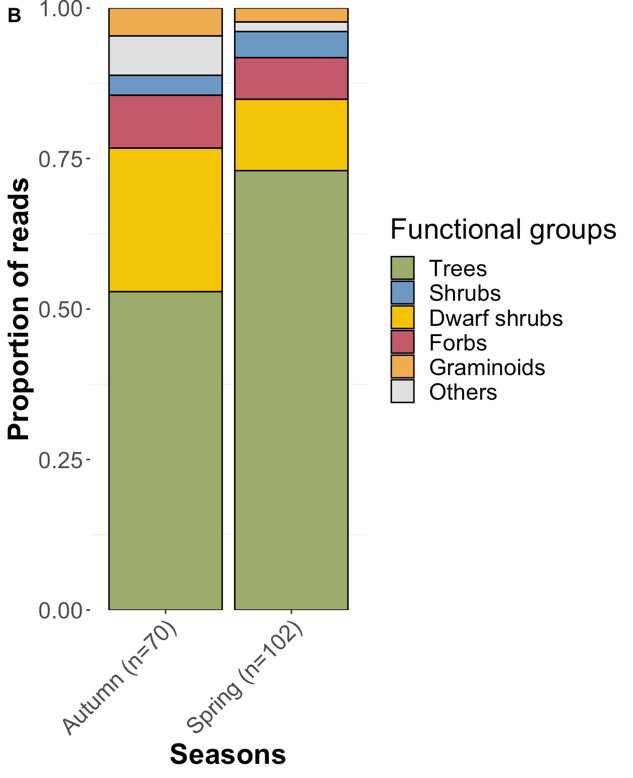

**Fig S2.3:** Dietary composition of plant functional groups retrieved from capercaillie faecal samples collected from **a)** both sexes and **b)** autumn and spring.

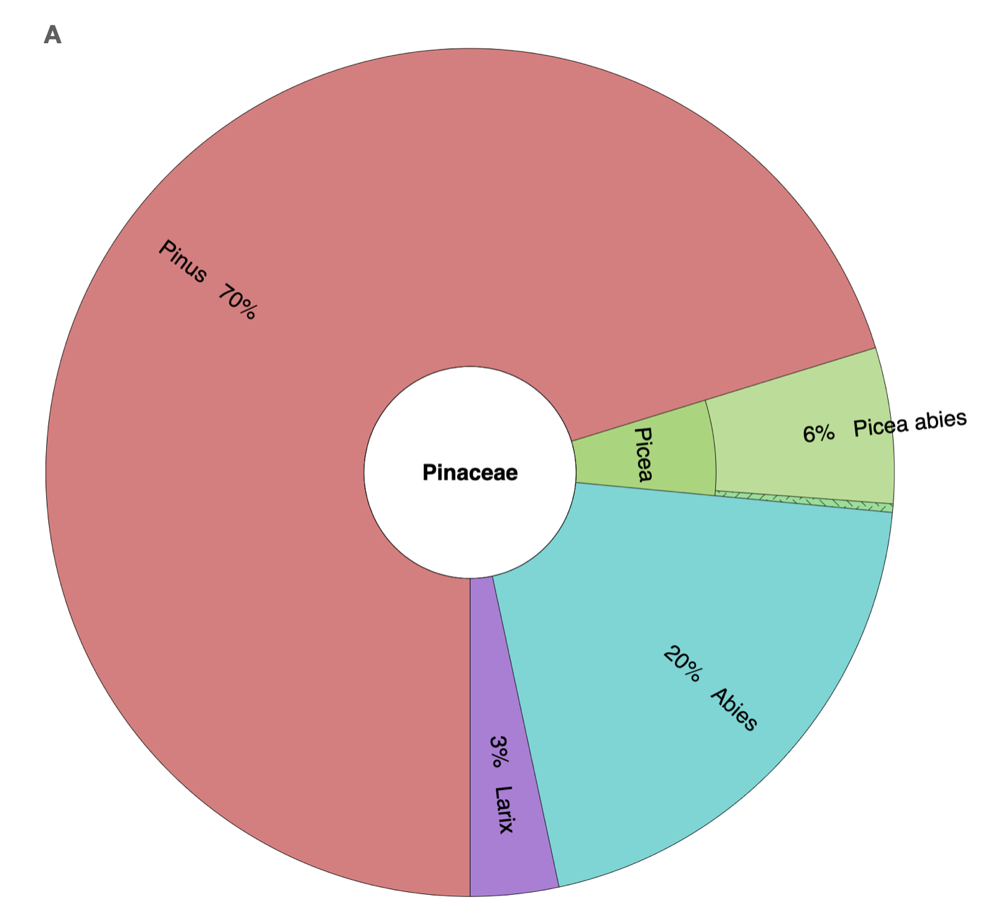

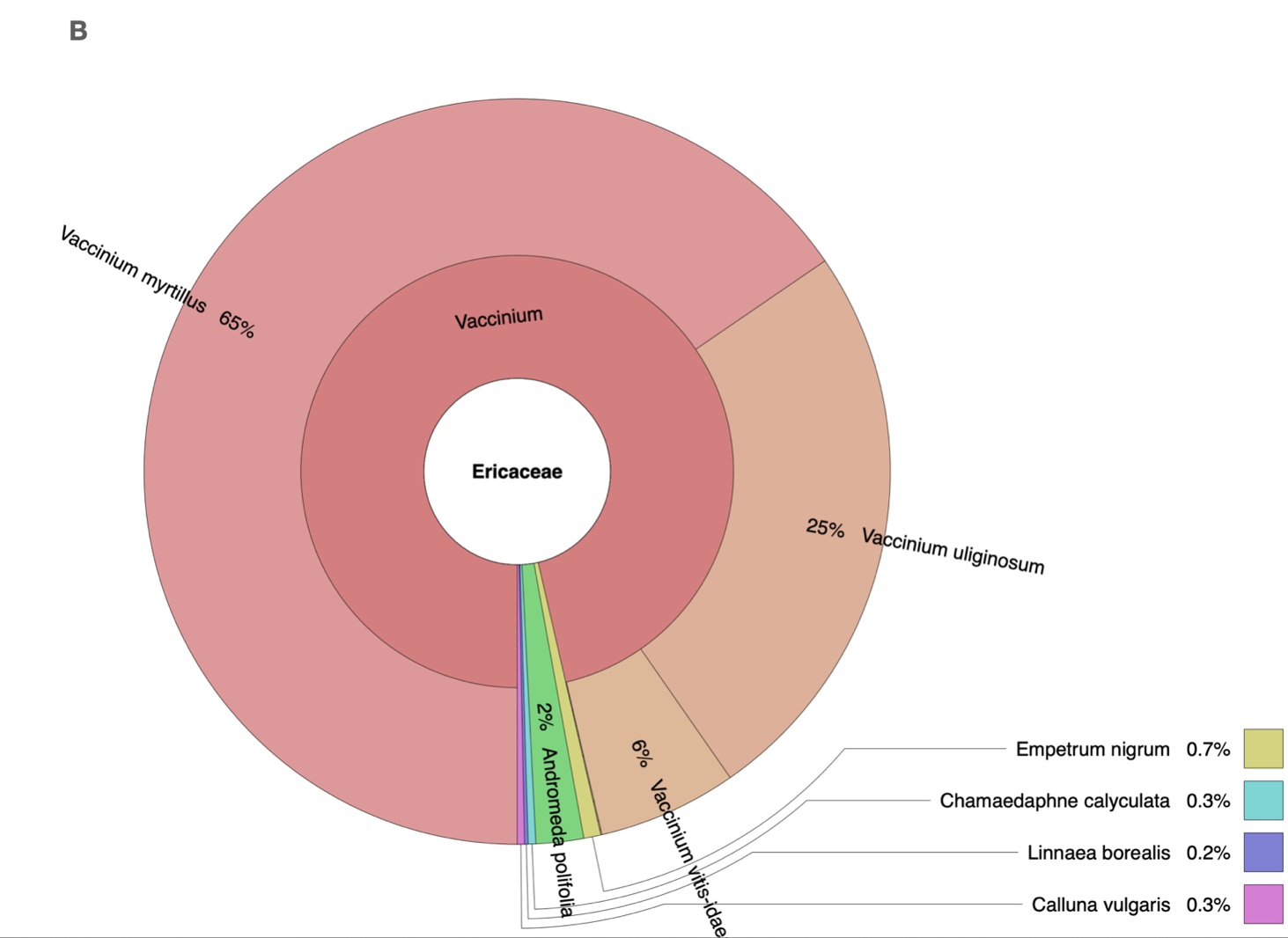

**Fig S2.4:** Krona chart showing the composition of plant taxa based on relative-read abundance (RRA) found in the families **a)** Pinaceae and, **b)** Ericaceae (Ondov et al., 2011).

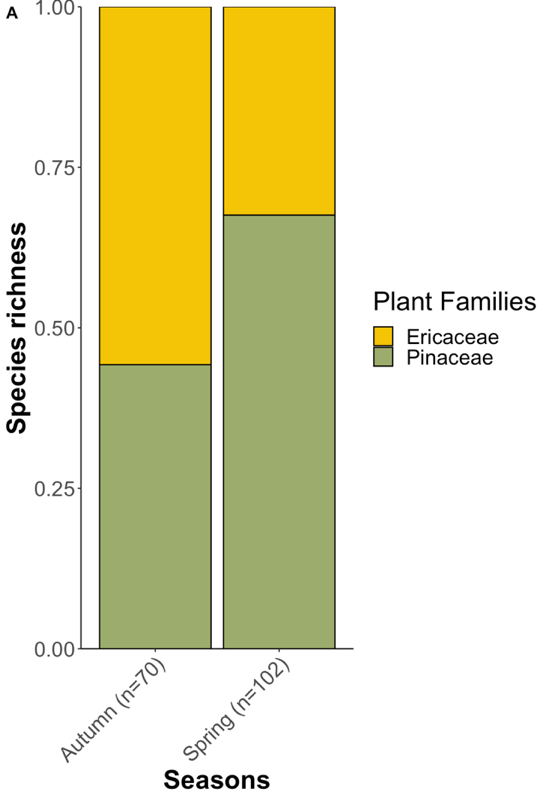

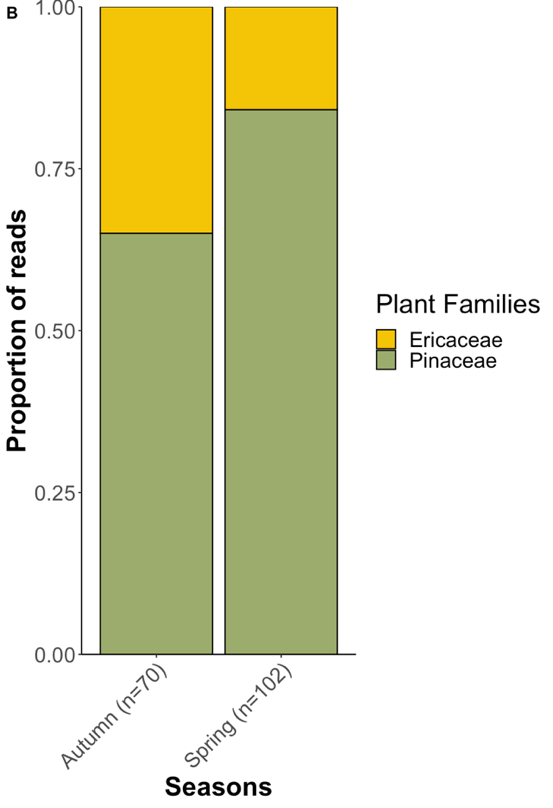

**Fig S2.5:** Seasonal shifts in Ericaceae and Pinaceae contents retrieved from capercaillie faecal samples based on **a)** species richness, and **b)** relative read abundance.

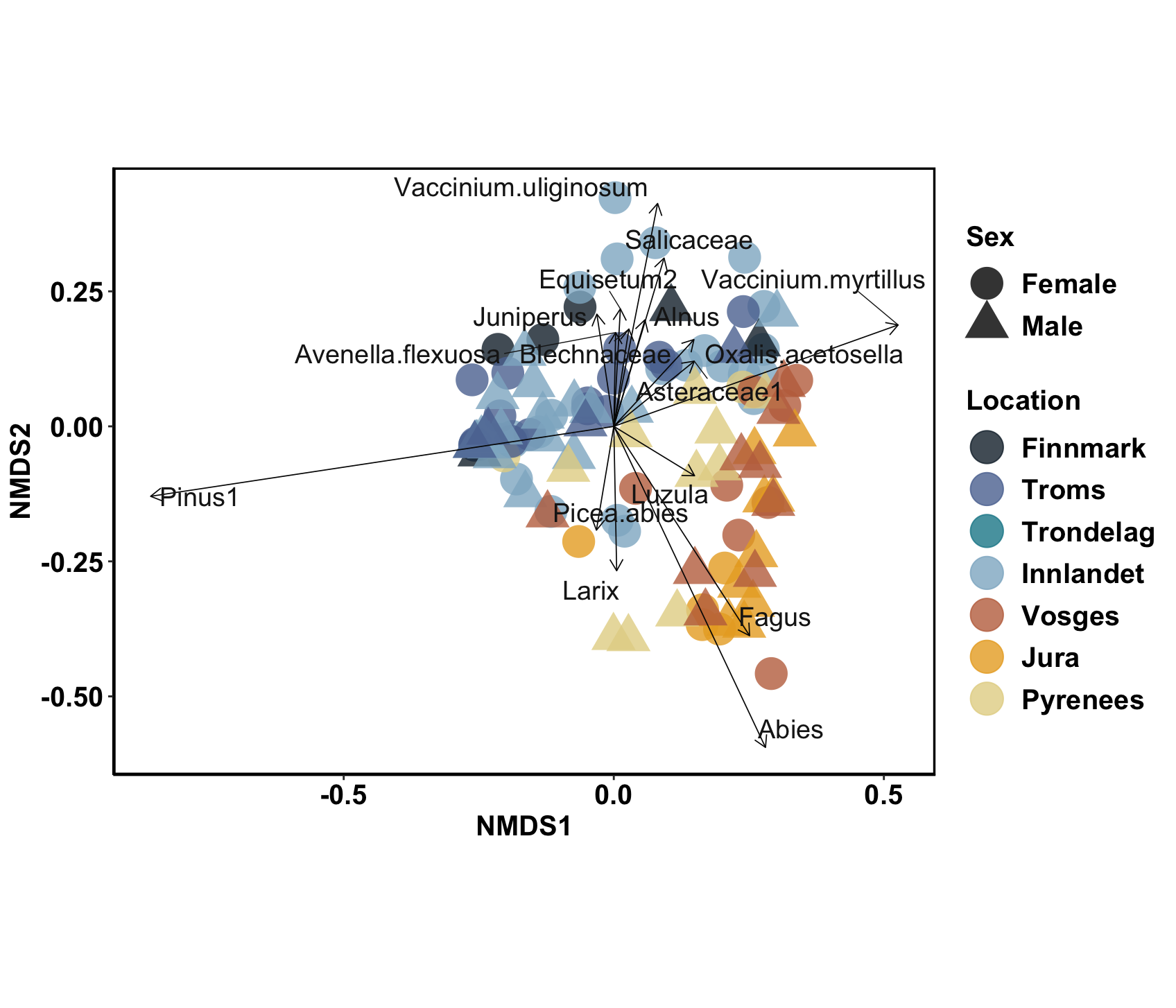

**Fig S2.6:** NMDS plots of RRA-based Bray-Curtis dissimilarity of capercaillie faecal samples collected from different locations (adonis F_5,86_ = 11.01, *r^2^* = 0.17, *p* = 0.001) and both sexes from each of these locations (adonis F_1,47_ = 2.77, *r^2^* = 0.04, *p* = 0.024) showing the plant taxa involved in driving distribution patterns. The stress level of 0.163 is under the acceptable value as suggested by Clark (1993) for an interpretable ordination.

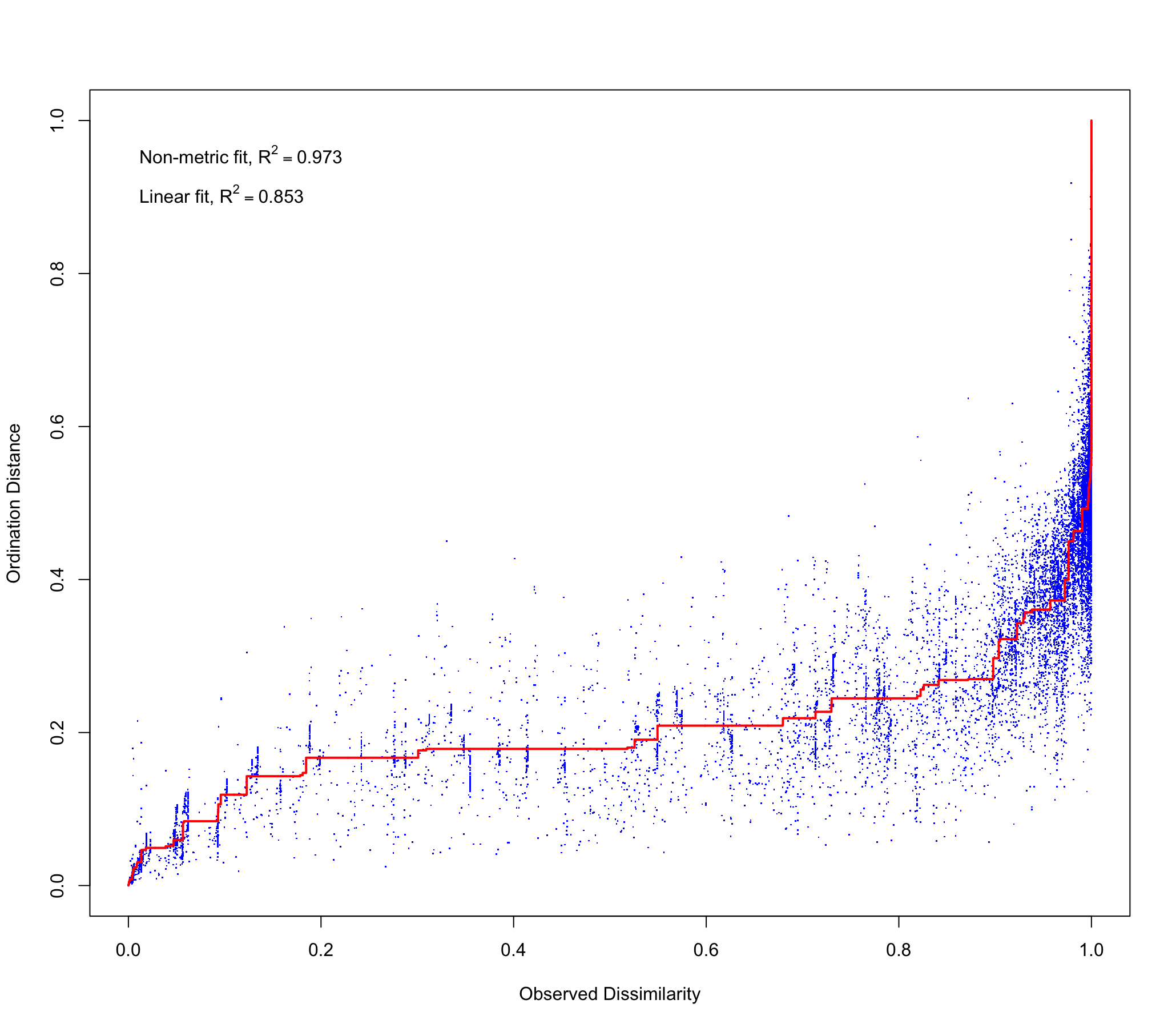

**Fig S2.7:** Shepard diagram showing correlation statistics (Non-metric fit R^2^ =0.973, Linear fit, R^2^ = 0.853) which indicates the fit between ordination distances and observed dissimilarities based on RRA data.
